# Voluntary alcohol consumption during distinct phases of adolescence differentially alters adult fear acquisition, extinction and renewal in male and female rats

**DOI:** 10.1101/2023.10.03.560757

**Authors:** J Alex Grizzell, Maryam Vanbaelinghem, Jessica Westerman, Michael P Saddoris

## Abstract

Alcohol use during adolescence coincides with elevated risks of stress-related impairment in adults, particularly via disrupted developmental trajectories of vulnerable corticolimbic and mesolimbic systems involved in fear processing. Prior work has investigated the impact of binge-like alcohol consumption on adult fear and stress, but less is known about whether voluntarily consumed alcohol imparts differential effects based on adolescence phases and biological sex. Here, adolescent male and female Long Evans rats were granted daily access to alcohol (15%) during either early (Early-EtOH; P25-45) or late adolescence (Late-EtOH; P45-55) using a modified drinking-in-the-dark design. Upon adulthood (P75-80), rats were exposed to a three-context (ABC) fear renewal procedure. We found that male and female Early-EtOH rats showed faster acquisition of fear but less freezing during early phases of extinction and throughout fear renewal. In the extinction period specifically, Early-EtOH rats showed normal levels of freezing in the presence of fear-associated cues, but abnormally low freezing immediately after cue offset, suggesting a key disruption in contextual processing and/or novelty seeking brought by early adolescent binge consumption. While the effects of alcohol were most pronounced in the Early-EtOH rats (particularly in females), Late-EtOH rats displayed some changes in fear behavior including slower fear acquisition, faster extinction, and reduced renewal compared with controls, but primarily in males. Our results suggest that early adolescence in males and females and, to a lesser extent, late adolescence in males is a particularly vulnerable period wherein alcohol use can promote stress-related dysfunction in adulthood. Furthermore, our results provide multiple bases for future research focused on developmental correlates of alcohol mediated disruption in the brain.

## Introduction

Alcohol is the most widely abused substance during adolescence, whether due to impulse, sensation seeking, social facilitation, or attempts to cope with the stress of this particular period of life (Bava and Tapert, 2010). Alcohol consumption rates remained high leading up to the COVID-19 pandemic, with over 4.2 million adolescents between the ages of 14 and 20 reporting binge consumption at least once in the month prior (Sicher et al., 2022). Alcohol exposure in adolescence can impart age-dependent, brain-related changes that present in adulthood as reduced resilience to stress-related neuropsychiatric dysfunction (Grant and Harford, 1995, Rohde et al., 2001, Bava and Tapert, 2010; Spear, 2018, Kyzar et al., 2016; Rohde et al., 1996).

Indeed, alcohol may exert particularly harmful effects on the brain during early-life development due to the unique vulnerability brain in this period. Early stages of life are comprised of sensitive developmental periods that establish the organizational trajectory of the brain as it structurally and functionally matures into adulthood. Neural development is marked by extensive neuronal proliferation in early childhood whereas adolescent periods are marked instead by intensive experience-dependent refinement of neural circuits, for example through experience-dependent reorganization, myelination, and pruning of neuronal ensembles (Gilmore et al., 2018). Thus, in resource-rich, socially supportive environments with minimal allostatic overload, decision-making and emotion-regulating brain regions typically organize in a manner that promotes adaptive, flexible responses to threatening situations in adulthood. Conversely, impoverished upbringing, poor familial/social support, and toxic environmental exposures can promote developmental trajectories that instead bias individuals towards inflexible and potentially maladaptive behaviors in response to stress and adversity later in life (Silbereis et al., 2016). Accordingly, adolescence is thought to be a particularly vulnerable period when exposure to stress, alcohol, or drugs of abuse disrupts brain development in ways that substantially increase the risk of developing stress-related disorders in adulthood, including posttraumatic stress disorder (PTSD), major depressive disorder, and substance use disorder (Fuhrmann et al., 2015; Sicher et al., 2022).

Adolescence can be broadly defined as the transitional period occurring between childhood and adulthood when numerous biological systems develop and mature, including the onset of puberty and sexual maturation (Spear, 2000). Adolescence in humans is generally thought to begin around 10-12 years and extend to 18-25 years (Spear, 2000, Crews et al., 2007), and efforts to determine development-dependent patterns of behavior often stratify adolescence further (*e.g.* early, middle, late phases). For example, while risk-taking occurs more often in adolescence than adulthood, impulsive behaviors tend to peak in early adolescence and decline linearly in a manner unrelated to pubertal maturation whereas sensation-seeking behaviors follow a pubertally linked, curvilinear pattern that peaks in middle adolescence (Steinberg et al., 2008). Indeed, pubertal onset typically begins during the early phase, albeit slightly later in males than females, and sexual maturity generally occurs during late adolescence (Dorn, 2006).

At the same time, the brain rapidly changes during these developmental phase-dependent periods as well. Within early adolescence, grey matter volumes peak in several corticolimbic regions and rapidly decline thereafter (Brenhouse and Andersen, 2011; Crews et al., 2007; Gass et al., 2014). However, white matter volumes continue to increase throughout the brain from early adolescence into adulthood, underscoring a linear progression of circuit development persisting into adulthood. As such, early adolescence may be a more sensitive developmental period for bottom-up mediated, socioemotional reactivity, motivation, and reward-processing whereas top-down control over emotion and decision-making may be at greater risk during late adolescence (Conrod, 2016; Spear, 2015, 2018). Disruptions during these phases (such as via alcohol or stress) could therefore produce distinct outcomes, with perturbations early in adolescence driving dysfunctional social/affective behaviors and substance abuse in adulthood, while such exposures in late adolescence primarily impacting cognitive tasks (Spear, 2015).

In particular, disruptions during adolescent developmental are known to alter corticolimbic trajectories that promote stress vulnerability later in life (Caballero et al., 2020). Indeed, deficits in fear conditioning, whether through enhanced fear learning, impaired extinction, or impaired recall in novel contexts, are key features underlying the negative consequences of many stress-related disorders, including PTSD (Maren and Holmes, 2016, Bouton et al., 2021). For example, patients with PTSD exhibit reduced context-appropriate renewal of fear compared with combat-experienced controls (Garfinkel et al., 2014). Animal models support these findings, especially in altered stress-induced anxiety, risk-taking, and memory (Fisher et al., 2017; Pajser et al., 2019, 2018). Furthermore, recent work shows that alcohol can have sex– and adolescent phase-dependent impacts on adult behavior in rats, where intermittent exposure to alcohol produced abnormal responses to both anxiety and reward. For example, alcohol in early adolescence was associated with increased non-social anxiety-like behavior in both adult males and females, while alcohol consumed in late adolescence only affected males (Varlinskaya et al., 2020). In contrast, late-but not early adolescent alcohol exposure induced inappropriate appetitive responses (Towner and Spear, 2021). Similarly, adult males with a history of early-mid adolescent alcohol exposure displayed deficits in contextual fear memory recall, while late-adolescent alcohol drinking instead impaired contextual fear extinction (Broadwater et al., 2013).

Despite these findings, much less is known about the impacts of voluntary alcohol use during developmentally-relevant phases of adolescence on subsequent adult behaviors, particularly while including female subjects and using complementary behavioral models to assess distinct components of fear learning. In this behavioral study, we assessed voluntary, binge-like alcohol consumption (or water-only controls) in either early (P25-45) or late (P45-55) adolescence in male and female Long Evans rats, then assessed fear-related learning in these animals later in their adulthood. Alcohol-mediated changes in fear acquisition, retention, extinction, and renewal were then tested using a Pavlovian, three-context, “ABC” fear renewal paradigm over three days (Jin and Maren, 2015). We predicted that adult rats with a history of early-adolescent alcohol would show deficits of cue-based fear learning and extinction whereas late-adolescent alcohol would impact context-appropriate freezing during fear renewal.

## Methods

### Subjects

Weanling Long Evans rats (n=27/sex; Envigo) were delivered on postnatal day 21 (P21) and housed 3/cage in same-sex, temperature– and humidity-controlled vivarium spaces on a 12/12 h light/dark cycle (lights on at 0700) with *ad libitum* access to food and water where they remained except during alcohol exposure periods and behavioral experimentation, as described below. All procedures followed guidelines for the National institute of Health and were approved by the University of Colorado, Boulder Institutional Animal Care and Use Committee.

### Adolescence: Alcohol Exposure

To encourage voluntary, binge-like alcohol consumption, we used a modified version of the Drinking-in-the-Dark (DID) paradigm (Thiele and Navarro, 2014). Rats had daily access to ethanol (EtOH) during either early-mid adolescence (P25-45, Early-EtOH; n=9/sex) or mid-late adolescence (P45-55; Late-EtOH; n=6/sex). Water-drinking subjects (n=12/sex) were used as age-matched controls. A schematic of the procedures used are shown in Fig 1A. For 4h at the start of the dark phase, rats were relocated within the vivarium to an experimental space and isolated in adjacent cages containing bedding, food pellets, and access single-housed housing environment where they were given access to a drip-resistant bottle that contained either water (for controls) or 15% EtOH in drinking water. Because a pilot study revealed that the DID paradigm did not impact water consumption after 2 days of habituation, a subset of water-drinking controls remained group-housed in their home cages during the DID to expedite throughput. To encourage greater consumption in the EtOH-assigned groups, the DID periods were extended to an average of 5h/day beginning at an approximate midpoint of the assigned drinking periods for both Early-EtOH (P35) and Late-EtOH groups (P49). Bottles were weighed before and after each assigned drinking period and animal weights were recorded immediately prior to the start of DID periods.

**Figure 1.**
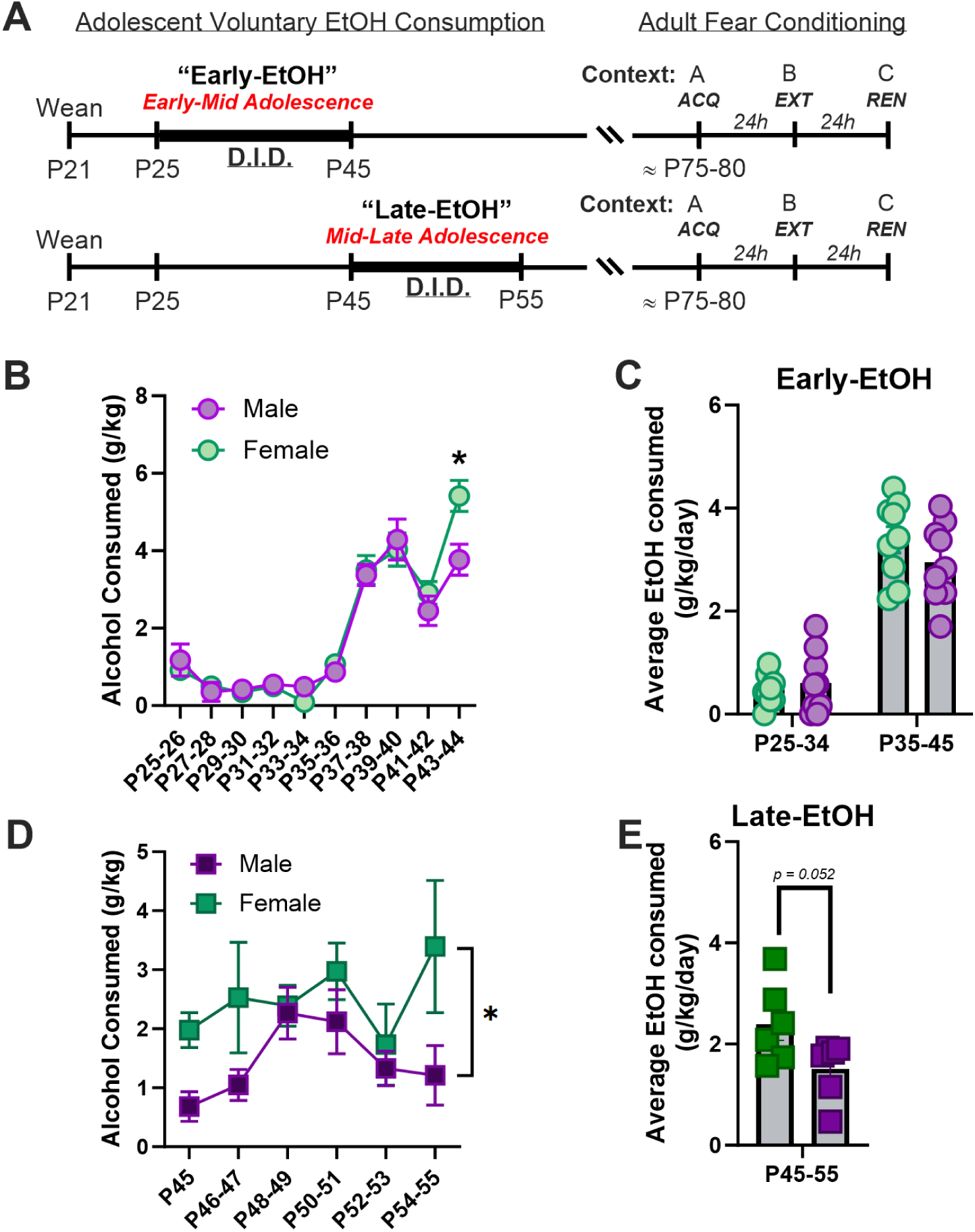
A. Schematic of the study design. Rats either consumed alcohol daily on a DID schedule early (top row) or late (bottom row) in adolescence. Upon adulthood, rats were trained in a three-context (ABC) fear renewal paradigm. B-C. Consumption (g/kg/day) for rats in the Early-EtOH group for two-day averages (B) or separated by early 4/h and later 5h periods (C). D-E. Consumption of alcohol for rats in the Late-EtOH shown in either 2-day averages (D) or overall for the entire adolescent period (E). *p<0.05, male vs female.

### Adulthood: Fear Acquisition, Extinction, and Renewal

After their respective drinking assignment periods, adolescents remained undisturbed in their home cages until adulthood (P75-80) at which time each rat was exposed to a three-context (“ABC”) fear renewal paradigm taking place over three consecutive days (Jin and Maren, 2015). Although contextual cues were changed daily, all testing was conducted within the first 4 hours of the light phase in the same dedicated procedure space with white and red-shifted fluorescent overhead lighting. Training occurred in four identical fear conditioning chambers (43 × 43 × 53 cm; Med Associates, St. Albans, VT) wherein inserts and other contextual stimuli were used to generate easily discriminable spaces. By default, these chambers were equipped with aluminum side walls, a transparent Plexiglas door and ceiling, a grid floor comprised of stainless steel rods placed 2 mm apart, and a removable tray below the floor grid (Bercum et al., 2021, 2023). In addition, the boxes could be illuminated from above using near-infrared light or a centrally located LED house light and were insulated from external stimuli via a sound-attenuating cabinet with a digital camera mounted on the interior of each cabinet’s door. The cabinets were also equipped with a superfluous fan for auditory ambiance and a side-mounted speaker external to conditioning chambers to deliver an auditory conditioned stimulus (CS). Visual CS was delivered via a white LED cue light (2 cm diameter) mounted on one side wall immediately above the auditory CS speaker port.

<u>Day 1 – Fear Acquisition (*Context A*)</u>: Rats were individually transported from the vivarium housing space to the conditioning room that was illuminated by white fluorescent lights via a white, 20-L bucket covered with green butcher paper. The ventilation fan and overhead LED housing light in each fear conditioning chamber remained ON, yet no additional modifications were introduced. Buckets and conditioning chambers were cleaned with 1% acetic acid. Fear acquisition training sessions consisted of a 3-min baseline exploration period followed by exposure to five cue (auditory tone [2kHz; 10s; 80dB] plus visual stimulus [white LED cue light]) pairings followed immediately by a footshock (1mA, 2s) and 60s inter-trial intervals (ITI).

<u>Day 2 – Fear Extinction (*Context B*)</u>: Rats were transported to the fear conditioning room that was illuminated by red-shifted fluorescent lights via a blue, 10-L bucket that was covered with a black, opaque Plexiglas lid. The ventilation fan and overhead LED housing light in each chamber remained OFF and a single custom-fit, curved, white, opaque Plexiglas insert (MedAssociates) was added to occlude the three aluminum sidewalls, yet without blocking audiovisual CS presentations. Buckets and chambers were cleaned with 70% EtOH. Extinction sessions began with a 3-min baseline period to assess activity and exploration, followed by 45 CS presentations (30s ITI).

<u>Day 3 – Fear Renewal (*Context C*)</u>: Rats were transported to the fear conditioning room that was again illuminated by red-shifted fluorescent lights via white, 20-L buckets with fresh bedding (approximately 3 cm deep) and a red Plexiglas lid that permitted some light exposure. The ventilation fan was turned ON and overhead LED housing light remained OFF. A custom-fit, flat, white, opaque Plexiglas insert (MedAssociates) was placed on the floor of the conditioning chamber, covering the grid floor. Boxes and buckets were thoroughly cleaned with Liquinox (Alconox, White Plains, NY). Renewal sessions began with a 3-min baseline period followed by five CS presentations (30s ITI).

### Data Analyses

Alcohol consumption was calculated for each animal by accounting for the EtOH concentration and animal weight (g/kg) for each DID drinking session. The expression of fear-related behavior data was collected using automated quantification of freezing with tracking software (VideoFreeze; MedAssociates) wherein digital images (collected at 30Hz) used frame-to-frame changes in pixel illumination to estimate movement. Freezing was defined as 30 consecutive frames (1 sec) where movement was subthreshold at or near zero, while freezing offset identified resumption in movement (>18 pixels/frame). While recent reports suggested that movement (e.g., darting) could differ between males and females (Gruene et al., 2015), neither automated motion detection nor coded observations by a trained observer revealed differences; as such, only freezing data are reported. Data are expressed as a percentage of time spent freezing (freeze onset to offset) within the defined periods of interest, which include baseline (BL), CS+ITI (Trial), and CS presentation alone (Cue). During extinction, Trial and Cue periods were averaged into blocks of five sequential epochs. Data were statistically analyzed using Student’s t-test and multifactorial ANOVA followed by *post hoc* analyses, where appropriate. BL was included for all Trial-based omnibus tests only. Because omnibus ANOVAs during each day of fear renewal revealed main effects/interactions of sex, separate analyses were conducted within males and females. Upon significance, predetermined (*a priori*) contrasts within epochs (e.g. male vs female, Early-EtOH and Late-EtOH vs Control, Early-EtOH vs Late-EtOH) were conducted using Fisher’s LSD, whereas all other contrasts were considered following Tukey’s HSD or Bonferroni’s correction, where described. Statistical analyses were conducted using Statistica (Dell) and graphed with Graphpad Prism.

## Results

### Alcohol Consumption during Adolescence

Rats in both the Early-EtOH (Fig. 1B-C) and Late-EtOH (Fig 1D-E) groups voluntarily consumed alcohol during DID. For the Early-EtOH rats, drinking increased across DID sessions, particularly when DID period extended from 4h/day (P25-34) to ∼5h/day (P35-45). Here, both Early-EtOH females and males significantly escalated consumption. Taking two-day averages, we found that there was no main effect of Sex, F_(1,16)_=0.65, p=0.43, though there was a main effect of Day, F_(9,144)_=75.51, p<0.0001, and an interaction between Sex X Day, F_(9,144)_=2.12, p=0.03. Post-hoc analysis of this interaction indicated that on the last two-day period (P44-45), female rats consumed more than males, (Tukey, p=0.012; Fig. 1B). Taking the overall average did not indicate any significant differences between male and females for either of the early days (t_(16)_ = 0.58, p=0.57) or late days (t_(16)_ = 1.21, p=0.25) of consumption (Fig. 1C).

Late-EtOH rats likewise consumed daily EtOH consumption from the phase (P45-48) to late phase (P49-55). Unlike the Early-EtOH cohorts, consumption during this later adolescent phase started at a higher level than the initial sessions of the Early-EtOH group, and as such, these Late-EtOH drinkers failed to show a significant increase over sessions (main effect of Day, F_(5,50)_=1.69, p=0.15). Likewise, there was no interaction between Sex X Day, F_(5,50)_=0.99, p=0.43. However, we found a modest but significant main effect of Sex, F_(1,10)_=5.82, p=0.04, though a simple main effects posthoc analysis did not reveal any reliable differences between males and females on any two-day average periods (all Tukey p>0.27; Fig 1D). These differences were nearly significant when taking the overall average drinking amounts for the Late-EtOH period, t_(10)_ = 2.21; p = 0.052 (Fig. 1E).

### Fear Acquisition during Adulthood

Upon adulthood (P75-80), rats were trained in a three-context (ABC) Fear Renewal paradigm. Here, we analyzed fear acquisition data (Day 1, Context A) in two key ways. First, we used the conventional approach in fear paradigms wherein freezing during the baseline period was included alongside each *Trial*, which comprised the percentage of time freezing during a CS cue period and the ITI period that followed (Figure 2A-C). Separately, we analyzed freezing that occurred during each CS (“Cue”) period only (Figure 2D-F). Following the conventional *Trial*-based approach, an omnibus multifactorial ANOVA of freezing (Fig. 2A) revealed significant main effects of Alcohol (F_(2, 48)_=3.24, p=0.048) and Trial (F_(5, 240)_=174.9, p<0.00001) with an trending interaction between the two (F_(10, 240)_=1.65, p=0.09). However, there was no main effect of sex, F_(1, 48)_=0.63, p=0.43. *Post hoc* analyses of the effects of Alcohol confirmed that Early-EtOH rats froze more than Late-EtOH (p=0.02) and a trending difference compared to water (p=0.09) counterparts.

**Figure 2.**
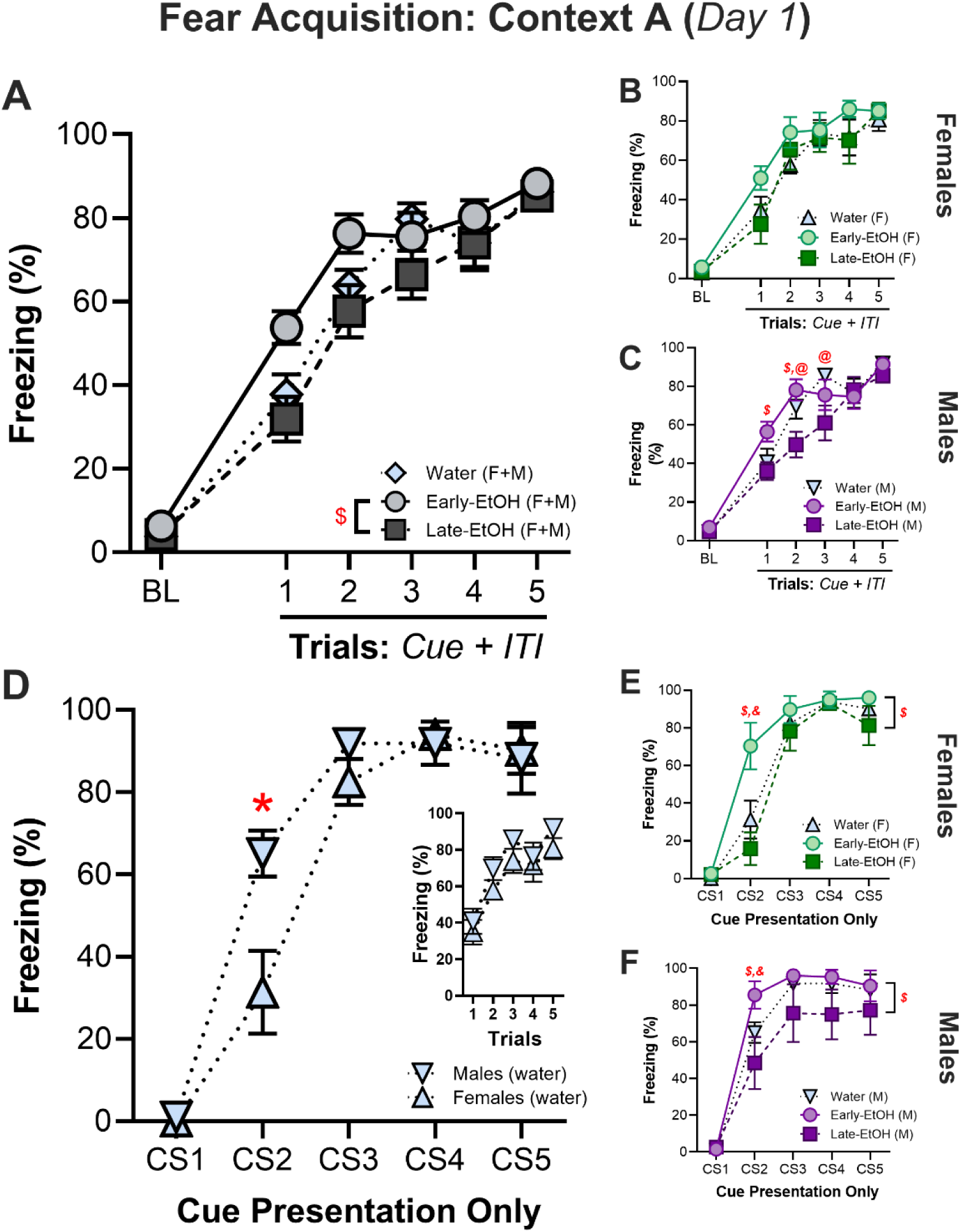
Adolescent alcohol consumption alter acquisition of fear in adulthood. A-C. Freezing during Trials (10s Cue+60s ITI). A. Omnibus test of fear acquisition indicates that Early-EtOH rats freeze more, particularly during early trials, than Late-EtOh and water controls. B. Female rats (all alcohol groups) and C. male rats across fear acquisition Trials. D-F. Data showing activity during the CS alone indicates emergent differences by sex. D. Comparison between male and female controls during fear acquisition. E. Female rats (all alcohol groups) and F. male rats during the CS presentations in fear acquisition. $p<0.05, Early-EtOH vs Late-EtOH; *p<0.05, male vs female; &p<0.05, Early-EtOH vs Water; @p<0.05, Late-EtOH vs Water.

To better understand this effect, we then separately investigated the effects of alcohol with the Cue+ITI Trial average for females and males. For females (Fig. 2B), there was no main effect of Alcohol (F_(2, 48)_=1.66, p=0.21) or interaction between Alcohol and Trial, F_(10, 120)_=0.75, p=0.67, but there was a main effect of Trial (F_(5, 120)_=79.05, p<0.00001; Fig. 2B). On the other hand, omnibus multifactorial ANOVA of freezing during the Cue alone revealed an additional interaction of Sex and Cue (F_(4, 192)_=6.27, p=<0.0001). In all cases, the most robust differences in freezing among groups were upon presentation of CS2 (*i.e.* the first cue presentation after the shock). Posthoc tests indicated that Early-EtOH subjects froze more than Late-EtOH and controls (all Trial and Cue contrasts, p<0.00001), Late-EtOH subjects froze less than controls (Cue: p<0.05; Trial: p=0.053), and males froze more than females (Cue: p<0.00001). Different patterns observed in males (Fig. 2C), which indicated that alcohol-mediating patterns in freezing during acquisition more readily emerged during Cue+ITI presentations. We saw a trend towards a main effect for Alcohol, F_(2, 24)_=2.8, p=0.08, and a main effect of Trial, F_(5, 120)_=99.9, p<0.0001, but unlike in females, also saw an interaction of Alcohol X Trial, F_(10, 120)_=1.98, p=0.04. This interaction was due to both an accelerated learning in the Early-EtOH and slowed acquisition in Late-EtOH. This manifested as Early-EtOH rats freezing more than the Late-EtOH (Trial 1, p=0.02; Trial 2, p=0.002) and more than Water controls (Trial 1, p=0.04) early in acquisition. Conversely, Late-EtOH rats showed reliably less freezing than Water controls on Trial 2 (p=0.02) and Trial 3 (p=0.004).

We then assessed freezing that occurred selectively during the Cue (Fig. 2D-F). While there were no detectable differences in Trial-based analyses (Fig. 2D, *insert*), we did not see any effect of Sex, F_(1, 48)_=1.8, p=0.019, but we did see significant interactions of Sex with other factors such as Trial X Sex, F_(5, 240)_=5.62, p<0.0001. This effect was due to greater freezing in males during CS2 (p<0.01; Fig. 2D). Given this (and that sex differences emerged throughout omnibus testing across our fear paradigm), male and female rats were also analyzed separately.

For the females during the Cue alone (Fig. 2E), there were main effects of Alcohol (F_(2, 24)_=3.82, p=0.036) and Trial (F_(5, 120)_=151.01, p<0.00001) with an interaction between them (F_(10, 120)_=3.09, p=0.002). Specifically, Early-EtOH females froze more than Late-EtOH (p=0.002) and controls (p=0.0004) during CS2. Across these analyses, differences were more robust during Cue-alone presentations than the averaged Cue+ITI, suggesting that adolescent alcohol-induced changes in fear acquisition among females preferentially impacts Cue-specific freezing and normalizes during ITI periods, especially early in the acquisition process.

For the males during the Cues alone (Fig. 2F), there was a main effect of Alcohol, F_(2, 24)_=3.43, p=0.049; Fig. 2F, and Trial, F_(5, 120)_=135.2, p<0.0001, but no interaction with Alcohol by Trial, F_(10, 120)_=1.29, p=0.24. *Post hoc* tests reveal that the main effect of Alcohol was due to Early-EtOH displaying overall more freezing than Late-EtOH (p=0.04). Furthermore, Late-EtOH males froze less than water on Trial CS2 (p=0.02) and than Early-EtOH on CS2 (p=0.0005) and CS3 (p=0.05).

Given that freezing during ITI periods differed, particularly among males, we also looked at responding in the 10-sec post-shock period (PostCue) for each cue of acquisition (*data not shown*). However, post-shock freezing did not appear to be affected by either sex or prior alcohol experience. There was no main effect of sex (F_(1, 48)_=0.39, p=0.56) nor any interactions of sex with other factors (Sex X PostCue: F_(4, 192)_=0.88, p=0.48; Sex X PostCue X Alcohol: F_(8, 192)_=1.05, p=0.40). Analyses of the effects of alcohol experience within sex likewise did not reveal any effects in either males or females (Females-Alcohol: F_(2, 24)_=1.27, p=0.29; Alcohol X PostCue: F_(8, 96)_=0.87, p=0.54; Males – Alcohol: F_(2, 24)_ = 2.37, p=0.12; Alcohol X PostCue: F_(8, 96)_=1.81, p=0.08).

### Fear Retention and Extinction

Twenty-four hours after fear acquisition, rats were tested for fear retention and extinction in a novel Context B via 45 trials (40s=10s cue+30s ITI); data were averaged into 5-Trial Blocks, except where noted. Note that, consistent with the analyses conducted for fear acquisition, we report data first for timepoints averaged across both the cue and the immediately-following post-cue ITI period (Fig 3A,C,D; left column), and again with the cue-presentation alone (Fig 3B,E,F; right column). First when we examined freezing during Cue+ITI with all sexes and ages analyzed together in an omnibus multifactorial ANOVA of Trial Blocks (Fig. 3A), there was no effect of alcohol generally, Alcohol (F_(2, 48)_=2.67, p=0.08), and a nearly-significant main effect of Sex (F_(1, 48)_=3.80, p=0.057), though there was a large main effect of Trial Block, F_(9, 432)_=49.95, p<0.0001. Despite these modest main effects of treatment, we found several robust interactions between factors including Alcohol X Trial Block, F_(18, 432)_=3.89, p<0.0001, Sex X Trial Block, F_(9, 432)_=1.97, p=0.04, and Alcohol X Sex X Alcohol, F_(18, 432)_=1.71, p=0.035. *Post hoc* investigations of these effects indicated less freezing overall in females than males (p=0.042) and the main effect of Alcohol was due to less freezing in the Early-EtOH than Water controls (p=0.03), but no differences between Early-EtOH and Late-EtOH (p=0.39) or Late-EtOH and Water (p=0.27). The Trial Block X Alcohol interaction was due to Early-EtOH exhibiting less freezing than both the Water controls and Late-EtOH during both the BL (both p<0.02) and the first trial block (both p<0.0001). However, later trials did not differ between these groups (all p>0.13), nor were there any differences between Late-EtOH and Water at any point (all p>0.10).

**Figure 3.**
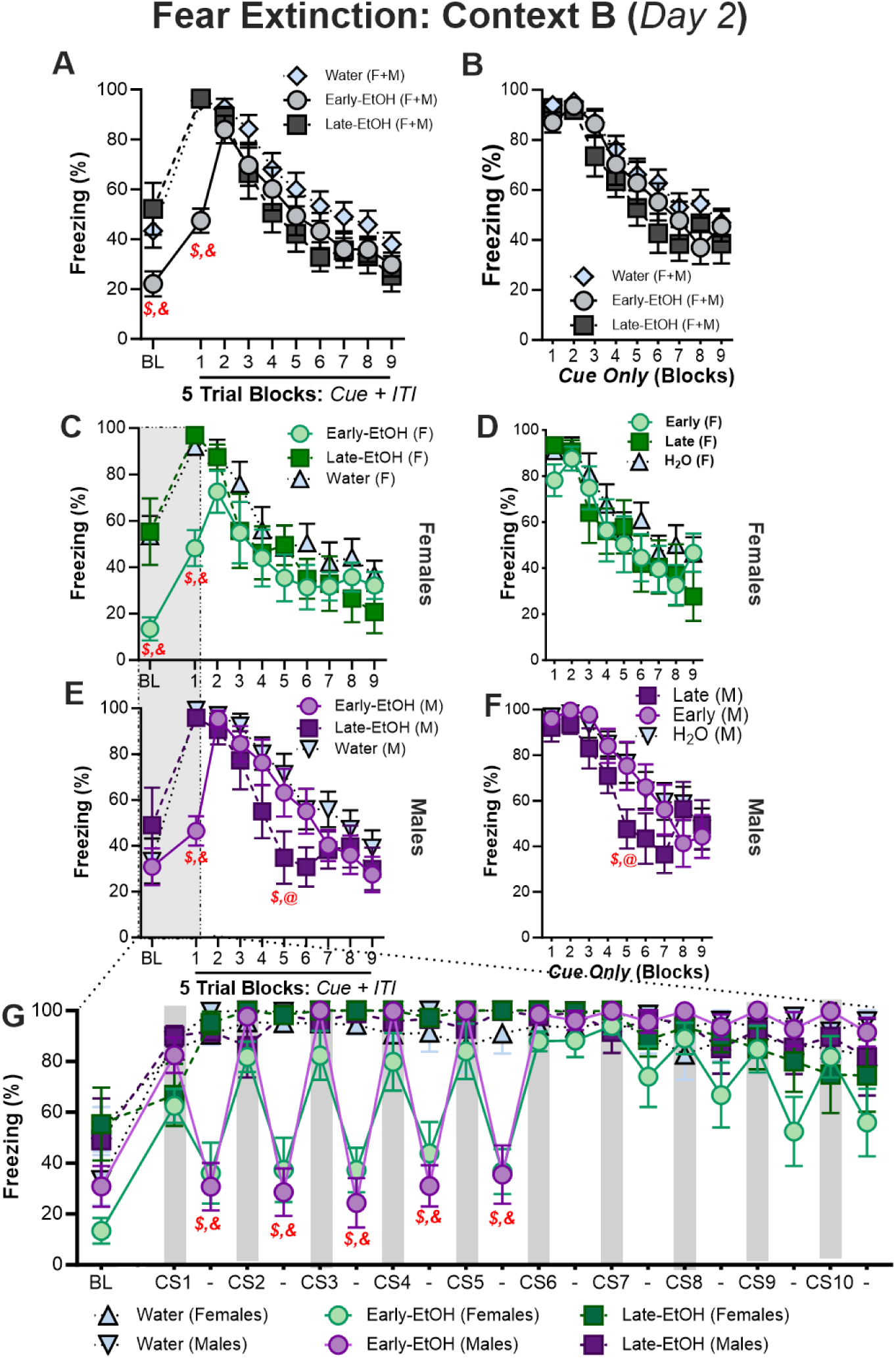
Adolescent alcohol differentially alters fear retention, extinction, and context discrimination. A. Freezing during baseline (BL) as well as Trials (10s Cue + 30s ITIs) averaged into blocks of 5 trials. B. Similar to A, but showing freezing only during the cue period (i.e., without the subsequent ITI). C-D. Female rats throughout fear extinction training (5-Trial Blocks) using the Cue+ITI average period (C) and the cue-only (D) periods. E-F. Male rats throughout fear extinction training (5-Trial Blocks) using the Cue+ITI average period (E) and the cue-only (F) periods.. G. BL (180s), CS (10s), and ITIs (30s) graphed chronologically for first 10 CS presentations of Extinction reveal distinct patterns of freezing in Early-EtOH rats only during early Cues but no other periods. $p<0.05, Early-EtOH vs Late-EtOH; *p<0.05, male vs female; &p<0.05, Early-EtOH vs Water; @p<0.05, Late-EtOH vs Water

Intriguingly, we discovered that these effects disappeared when we looked at freezing during just the Cue period (i.e., without the post-cue ITI included), including no effect of Alcohol, F_(2, 48)_=1.29, p=0.29; Fig. 3B) and only a trend towards a main effect of Sex (F_(1, 48)_=3.77, p<0.058). While there was no interaction of Trial Block with Sex, F_(9, 432)_=1.32, p=0.22, or Trial Block X Sex X Alcohol, F_(18, 432)_=1.14, p=0.31, we did see a significant interaction of Trial Block X Alcohol, F_(18, 432)_=1.90, p=0.015. However, this was due exclusively due to decreased freezing in the Early-EtOH animals versus the Late-EtOH (p=0.002), and versus Water (p=0.009), with no differences beween Late-EtOH and Water (p=0.33); there were no differences in freezing during any of the subsequent cue presentations (all p>0.18).

When we separately assessed freezing for the Cue+ITI periods by sex, females (Fig. 3C-D) showed a no main effect of Alcohol F_(2, 24)_=1.69, p=0.21, but there was a significant interaction between Alcohol and Trial Block (F_(18, 216)_=2.22, p=0.004; Fig. 3C). Indeed, these Cue+ITI differences were carried by the fact that Early-EtOH females froze robustly less during earliest (retention) periods of extinction (Fig. 3C) than Late-EtOH (BL, p=0.002; Trial Block 1, p=0.001) and than water controls (BL, p=0.006; Trial Block 1, p=0.001), while Late-EtOH and Water never differed during these points (both p>0.63), and indeed no differences between any groups for a given trial block were found from Trial Block 2 onward (Fig. 3C). Notably, these interactions were diminished when assessing freezing only during the Cue period, where there was no main effect of Alcohol, F_(2, 24)_=1.00, p=0.39, but a modest interaction between Alcohol and Trial Block (F_(18, 216)_=1.74, p=0.04; Fig. 3D). However, this latter interaction was due again to differences during the Early-EtOH and the other groups during the pre-cue BL (both p<0.005) but no differences during any of the Trial Blocks where cues were presented (all p<0.15).

We observed slightly different effects in males (Fig. 3E-F). During the Cues+ITI, there was no main effect of Alcohol, F_(2, 24)_=1.14, p=0.37, though there was a robust interaction between Trial Block and Alcohol, F_(18, 216)_=3.35, p=0.0003 (Fig. 3E). Post hoc analysis showed that this interaction was due to two distinct effects in the extinction sequence. First, early n extinction, Early-EtOH rats significantly reduced freezing in the first Trial block compared to both Water (p<0.0001) and to Late-EtOH (p=0.0004), while Late-EtOH and Water did not differ in this block (p=0.80). Thus, unlike in females, there was no effect of alcohol in the pre-cue BL period. A second effect emerged later in extinction where freezing in the Late-EtOH rats rapidly decreased at a rate faster than the other groups; by Trial Block 5, Late-EtOH rats were freezing significantly less than both Early-EtOH (p=0.03) and Water (p=0.01; Figure 3E).

Looking selectivey at the Cue-only period in the males, we again found no effect of Alcohol, F_(2, 24)_=0.76, p=0.48, or an interaction between Trial Block X Alcohol: F_(18, 216)_=1.34, p=0.16; Fig. 3F). Planned post hoc comparisons indicated that, while the effect of accelerated extinction in the Late-EtOH group compared to other groups on Trial Block 5 was still observed (vs Water, p=0.02; vs Early-EtOH, p=0.03), there was no longer an differences between groups on the first Trial Block as had been seen when using the Cue+ITI average. Thus, Late-EtOH rats appeared to more rapidly extinguish cued fear memories than controls and Early-EtOH counterparts.

Given the disparity between Blocked Cue– and Trial-based analyses, we assessed changes in freezing during first 10 trials by splitting the CS and ITI periods (Fig. 3G). We found a trending effect of Sex (F_(1, 48)_=3.26, p=0.08) and a main effect of Alcohol (F_(12, 48)_=12.9, p<0.0001) as well as interactions between Alcohol and all other factors, including a 3-way interaction between Alcohol, Sex, and BL/CS/ITI (F_(38, 912)_=1.47, p<0.05). Bonferroni-corrected comparisons indicated that both female and male Early-EtOH groups froze significantly less than controls and Late-EtOH during each of the first five ITI periods (all p<0.01), but not adjacent Cue periods. Notably, we did not detect differences between Early-EtOH males and females at any point in this analysis.

### Fear Renewal

Twenty-four hours later, rats received five CS presentations in Context C followed by 30s ITIs in a fear renewal test, which grants the opportunity to assess the degree to which animals show an adaptive, context-appropriate relapse of fear when presented with the potentially threatening CS in a novel context (Fig. 4). In an ominbus ANOVA that included both male and female subjects to assess freezing during the Cue+ITI period, we found a main effect of Alcohol, F_(2, 48)_=6.471, p=0.003, and a main effect of Trial, F_(5, 240)_=49.30, p<0.0001, though the main effect of Sex did not reach significance, F_(1, 48)_=2.76, p=0.10. *Post hoc* analyses of the main effect of Alcohol confirmed that Early-EtOH rats displayed less freezing than Water controls (p=0.001) and a Late-EtOH rats (p=0.03), while Late-EtOH did not differ from controls (p=0.50; Fig. 4A). Surprisingly, a planned *post hoc* comparison of the main effect of Sex indicated a small but significant difference between males and females (p=0.04 Fig. 4B). However, we did not see any interactions between Sex with any other factors (Sex X Alcohol, F_(2, 48)_=1.051, p=0.36; Sex X Trial, F_(5, 240)_=1.01, p=0.41; Sex X Alcohol X Trial, F_(10, 240)_=0.84, p=0.59).

**Figure 4.**
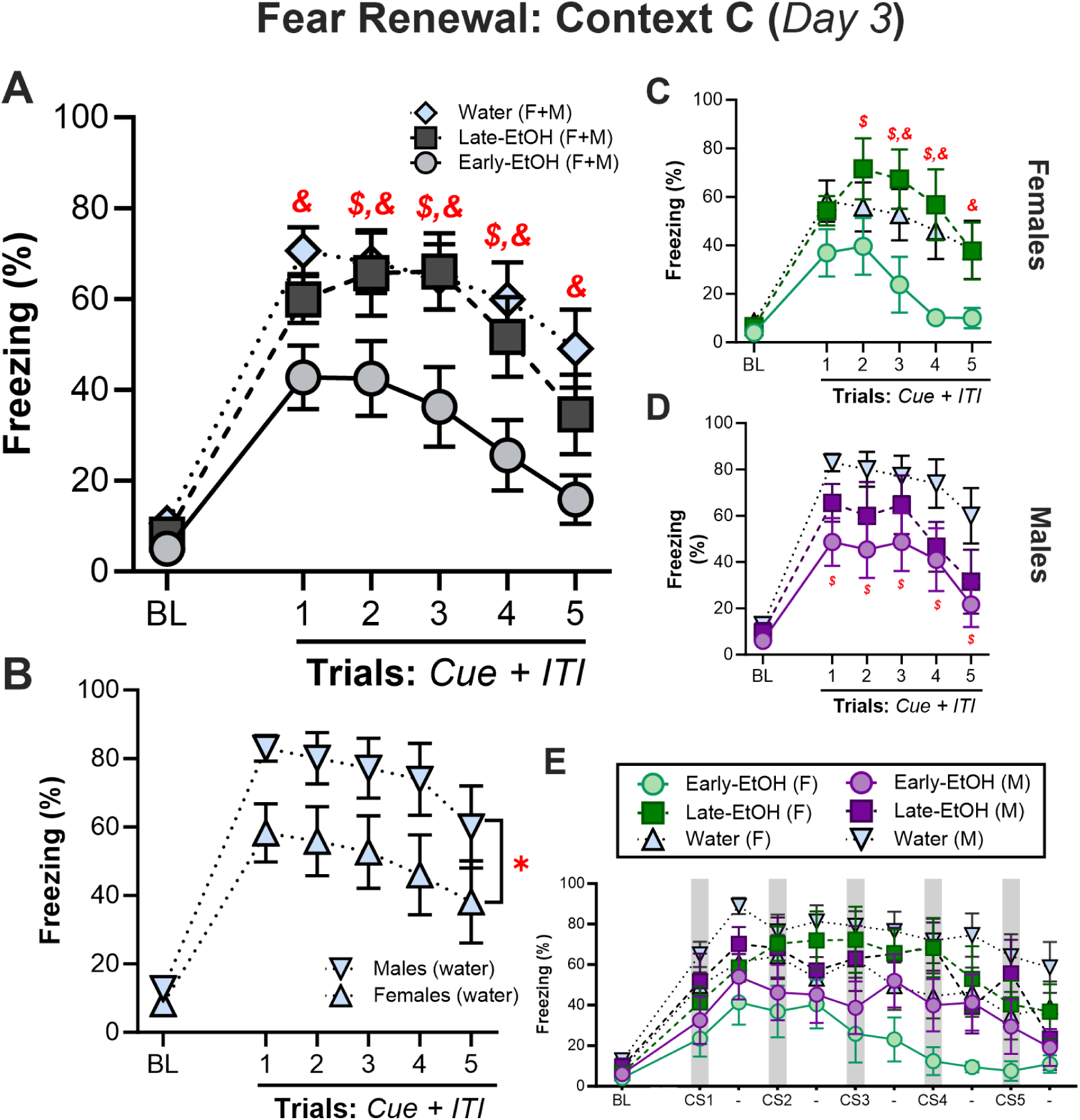
Adolescent alcohol impacts fear renewal in adulthood in a sex– and development-dependent manner. A. Omnibus test of freezing during fear renewal trials (10s Cue + 30s ITI) in context C reveal that Early-EtOH rats show substantially less context-appropriate renewal of fear. B. Male and female controls confirm inherent sex-differences in expression of fear renewal. C. Female rats and D. male rats across fear renewal test show sex-dependent differences in freezing among Late-EtOH rats. E. Trial-by-trial analysis showing freezing during both the 10s cue (shaded bars) and 30s ITI periods for all groups and treatments. $p<0.05, Early-EtOH vs Late-EtOH; &p<0.05, Early-EtOH vs Water; @Late-EtOH vs Water

We then separately investigated each sex to assess the effects of alcohol in fear renewal. In females (Fig. 4C), we found a trend toward a main effect of Alcohol (Trial: F_(2, 24)_=3.26, p=0.055) and a trend toward an interaction of Trials X Alcohol (F_(10, 120)_=1.73, p=0.08). Planned *post hoc* analyses of the effect of Alcohol confirm that Early-EtOH females froze less than controls (p=0.043) and Late-EtOH females (p=0.03), while there were no observed differences between Late-EtOH and Water (p=0.63). *Post hoc* comparisons to explore the interaction indicated significant decreases in freezing for Early-EtOH compared to Water on trial 3 (p=0.03), trial 4 (p=0.008), and trial 5 (p=0.04), but not during the BL or earlier cues. Similarly, Early-EtOH showed reliably less freezing than Late-EtOH on trial 2 (p=0.04), trial 3 (p=0.007), and trial 4 (p=0.004), but not the BL or trials 1 or 5. No differences between Water and Late-EtOH were observed (all p>0.29).

In males however, we saw slightly different patterns (Fig. 4D). Like females, there were main effects of Alcohol, F_(2, 24)_=4.30, p=0.025, and Trial, F_(5, 120)_=27.31, p<0.0001, though we did not observe an interaction between Alcohol X Trial, F_(10, 120)_=0.96, p=0.48. For the effect of Alcohol, Early-EtOH males froze less than Water controls (p=0.008), though not than the Late-EtOH group (p=0.37). There was no reliable difference between Late-EtOH and Water overall (p=0.13). Planned post hoc comparisons indicated that Early-EtOH rats froze less than Water controls on every trial except the BL (cue trials 1-5, all p<0.04), while no other pairwise comparisons between alcohol group on given trials differed from each other.

Finally, to determine whether the cue-specific freezing pattern we observed in Early-EtOH rats during extinction followed into renewal, we separated BL, CS, and ITIs periods. Although there was a main effect of Alcohol (F_(2, 48)_=7.02, p<0.01; Fig. 4E) driven by reduced freezing in Early-EtOH than controls (p<0.01) and Late-EtOH (p<0.05), there were no interactions of Alcohol or Sex with any other factors (all p values for interactions, >0.35).

## Discussion

Deficits in fear learning are common among many suffering from trauma– and stress-related disorders, including PTSD, depression, and anxiety (Maren and Holmes, 2016; Maren et al., 2013), and it is well-established that various forms of adversity during sensitive developmental periods are time-dependent risk factors for these adult disorders. Here, we assessed sex– and developmental phase-dependent effects of voluntary, binge ethanol consumption during adolescence on fear-associated behaviors in adulthood. In general, we found that sex and adolescent phase differentially impact alcohol-mediated fear learning. Regardless of sex, adult rats with a history of repeated alcohol consumption during early adolescence displayed wide-ranging changes in fear memory that may suggest dysfunction, including accelerated fear acquisition, attenuated fear retention, and decreased context-appropriate fear renewal. On the other hand, later alcohol experience produced opposing and sex-dependent changes in fear learning, suggesting Late-EtOH experience impacts males to a greater extent than females and may slow fear acquisition, speed extinction, and impede fear renewal. It is noteworthy that we also found sex-dependent differences in fear learning among water-drinking controls, particularly indicating differences in fear acquisition and renewal in adult Long Evans rats. Also, there were marginal sex-related effects during adolescence wherein females tended to consume more alcohol than males as they aged, which is consistent with previous studies (e.g., Amodeo et al., 2018; Szumlinski et al., 2019).

### Alcohol Consumed in Early Adolescence Increases Freezing During Acquisition

Early-EtOH male and female rats showed accelerated acquisition of associative fear, while in contrast, Late-EtOH either decreased fear learning (males) or had no effect (females) relative to controls. Notably, these alcohol-associated changes were most pronounced during the cue alone. These observations suggest, that adolescent alcohol exposure altered primarily the learned response to stimuli (i.e., during the CS), particularly early in acquisition, rather than to the trials (Cue+ITI period) as a whole. Second, Early-EtOH rats of both sexes appeared accelerate associative freezing behaviors. This indicates that alcohol experience during early adolescence may sensitize fear responses to cues during learning (Susskind et al., 2008), although some studies have not shown these differences (Broadwater and Spear, 2013). However, studies failing to show effects of early-alcohol-potentiated learning used only male Sprague Dawley rats and an auditory CS. Our use of a more salient audiovisual cue and both sexes may have better revealed changes in associative learning process in Early-EtOH rats.

We hypothesize potential mechanisms that could produce these changes. The basolateral amygdala (BLA) has been shown to play an essential role in assigning associative value to sensory inputs associated with salient outcomes, and therefrom permitting the expression of the conditioned fear (e.g., Johansen et al., 2011). In rats of both sexes, the lengths, complexities, and densities of BLA neurons increase substantially between postnatal day 20 (P20) to P35, reaching adult levels soon thereafter, though these effects are also related to onset of puberty (Koss et al., 2014; Spear, 2000). Still, most BLA cellular refinement processes do not occur until late adolescence (Rubinow and Juraska, 2009). As such, one interpretation of our acquisition data is that alcohol experience produces development-dependent disruptions of BLA maturation and processing. Indeed, as males mature into puberty slightly later than females, this prediction would be consistent with males remaining more vulnerable to alcohol exposure later than females.

Alternatively, these data may implicate alcohol-mediated changes in dopamine function and reflect recent findings that fear learning and discrimination can be accelerated by phasic increases in dopamine during conditioning. Mesolimbic projections from ventral tegmental area (VTA) to nucleus accumbens (Saddoris et al., 2018, 2015), and/or mesoamygdalar projections from VTA to BLA could potentiate acquisition of aversive learning and enhance discrimination between different threats (Jo et al., 2018; Tang et al., 2020). Accordingly, accelerated fear acquisition in the Early-EtOH group could be linked to better learning in limbic regions, including BLA.

### Fear Extinction Is Differentially Modulated by Timing of Adolescent Alcohol Experience

One of the most striking set of observations in our study was seen in the earliest phases of the fear extinction session, wherein retention of fear can be assessed. Despite showing rapid acquisition of fear the day before, Early-EtOH rats showed less freezing to the novel context in the baseline (pre-cue) period as well as the first block of extinction trials.

Several potential interpretations for these observations are worth exploration. First, alcohol during early adolescence could have impeded fear memory consolidation after Day 1 training thus resulting in reduced fear recall on Day 2. However, this appears unlikely given that freezing during cue-presentations reached maximal levels early on and did not differ among rats of any group in our study. Furthermore, reduced cue-elicited freezing among Early-EtOH rats was selective to ITIs periods, but otherwise normal (and high) during cues. Furthermore, this effect was transient and disappeared by the second block of trials.

An alternative interpretation is that water-drinking controls and Late-EtOH rats better detected common contextual elements across Days 1 and 2. Consistent with this, work has shown that early – but not late – adolescent alcohol exposure can induce deficits in contextual fear recall in adulthood (Broadwater and Spear, 2013). Indeed, earlier onsets of alcohol exposure in adolescence elicit greater electrophysiological alterations of hippocampal activity in adulthood (Slawecki et al., 2001), which could account for failures in contextual fear memory (Spear, 2015). However, this interpretation may only be true for freezing during the pre-cue baseline period as it does not fully explain why Early-EtOH rats increased freezing during the ITI as trials progressed.

We posit another potential interpretation: early adolescent alcohol exposure could alter the sensitivity of contextual information and/or heightens the rewarding nature of novelty. Early-EtOH rats may more readily engage in elevated risk-taking as they over-valued a desire to explore the novel Context B, despite it being an ambiguously threatening context due to cue presentation. Moreover, as rats explored Context B and the novelty subsides (*i.e.* Trial Block 2), circuit-level competition for fear-motivated behaviors may have ultimately driven the observed freezing response that then statistically match those of controls. Despite the aversive nature of Pavlovian fear conditioning, animals concurrently process contextual information in the background (Phillips and LeDoux, 1994). In this way, learning in novel situations can adaptively facilitate contextual discrimination. However, passive fear responses like freezing generally suppress activity necessary to evaluate novelty of contexts.

Little research has been dedicated to determining whether phase-dependent adolescent alcohol exposure drives novelty seeking in adulthood. Male Wistar rats with alcohol exposure throughout adolescence show indices of behavioral disinhibition (Desikan et al., 2014) and risky, impulsive choices (Boutros et al., 2014). In contrast, Kim et al. (2019) showed that early-mid adolescent exposure in male and female Sprague-Dawley rats did not alter novel object exploration in adulthood. However, novelty seeking paradigms in rodents (*e.g.* novel object vs. novel context) are often poorly correlated, even within animals, thus underscoring the notion that novelty seeking is a multifaceted construct (Bardo et al., 1996; Pawlak et al., 2008).

Furthermore, animals selectively bred to express high novelty seeking show relevant phenotypic traits such as impulsivity and compulsive behaviors (Flagel et al., 2014; Stead et al., 2006). Novelty seeking is thought to be mediated primarily by midbrain dopamine system and interpeduncular nucleus; for example, stimulating dopamine increases novelty seeking (Molas et al., 2017) while also decreasing anxiety-like behavior in open arm mazes (DeGroot et al., 2020).

We therefore hypothesize that Early-EtOH rats’ delayed expression of freezing during extinction could be due to trial-dependent increases in dopamine reactivity, which may have initially driven context exploration during the ITI when the immediate threats (cues) were absent. Supporting this interpretation, alcohol induces a more substantial shift in the developmental trajectory of early-mid adolescent rats than in late adolescence and adulthood, therein biasing greater basal dopamine levels in the nucleus accumbens after a history or repeated alcohol exposure (Philpot et al., 2009). Alcohol during mid-adolescence also potentiates stimulus-evoked phasic dopamine as well as incentive salience in conditioned approach (Spoelder et al., 2015). Dopamine release also enhances learning about contexts when stimulated in the hippocampus (Tsetsenis et al., 2021) and cues when stimulated in the BLA (Jo et al., 2018). Future studies should identify whether phases of adolescence differentially mediate alcohol-dependent impacts of dopamine function in limbic targets like hippocampus and amygdala during cue and context fear learning.

Despite alcohol-related differences very early in extinction, rats subsequently displayed similar patterns of freezing for the rest of the extinction session, although Late-EtOH males displayed more rapid extinction when compared to same-sex counterparts. This effect, as in initial fear acquisition (day 1) appears to be selective to the cue presentations, suggesting altered processing of associative information, and was not seen in females. Thus, these data suggest that these effect are linked to delayed pubertal onset in males producing unique vulnerabilities to alcohol later than in females.

### Adolescent alcohol’s effects on fear extinction and renewal in adulthood

Fear renewal paradigms are designed to assess the relapse of fear when a previously-extinguished fear memory is presented in a novel context Our study shows that adolescent alcohol exposure disrupts fear renewal in sex– and phase-dependent ways. Specifically, Early-EtOH rats, regardless of sex, showed reduced levels of freezing compared to controls throughout renewal testing, while Late-EtOH experience produced opposing effects in males (decreased renewal) and females (increased renewal). However, it is often adaptive for threat-related contingencies to be accessible, especially when navigating new environments posing unknown threats. Thus, the reduced fear seen in the alcohol-experienced rats in the renewal session points to impairments in prior extinction and recall in this new context.

Renewal paradigms have illuminated complementary neural circuits underlying fear extinction and renewal, respetively. For example, the mPFC plays critical roles in gating fear expression particularly in extinction (Maren and Holmes, 2016, Bouton et al., 2021). While prelimbic (PL) mPFC afferents to BLA promote conditioned fear expression, competing mPFC-BLA projections from the adjacent infralimbic (IL) cortex are critical for driving extinction (Maren and Holmes, 2016). However in fear renewal scenarios, the BLA receives substantial context-dependent signals from both the PL and ventral hippocampus (Orsini et al., 2011), which also projects to the mPFC (Jin and Maren, 2015; Wang et al., 2016). Indeed, studies in humans confirm that hippocampal activation was greater in novel rather than previously extinguished contexts (Hermann et al., 2016; Zabik et al., 2023), while patients with PTSD counterintuitively showed less fear relapse, a reduced skin conductance response, and less activity of the amygdala, mPFC, and hippocampus in a fear renewal context when compared with combat controls (Garfinkel et al., 2014). Thus, consistent with the reduced fear renewal in our study, pathologies in fear and anxiety (such as PTSD) may not necessarily be a loss of inhibitory regulation *per se*, but a loss of context-appropriate engagement of inhibitory mechanisms over fear and fear expression (Maren et al., 2013; Maren and Holmes, 2016).

Consistent with this, we hypothesize potential mechanisms may involve appropriate excitatory/inhibitory balance in fear-related limbic regions such as the mPFC, including the function of inhibitory interneurons (e.g., parvalbumin [PV+]) and mesocortical dopamine activity. For example, footshock activates mPFC PV+ interneurons (particularly in IL) in both males and females yet experimental PV+ manipulations drive fear extinction deficits in a sex-dependent manner (Binette et al., 2023). Furthermore, fear renewal requires PV+ interneurons in the IL of the mPFC, which are preferentially targeted by ventral hippocampal projections to promote fear expression in a feed-forward inhibitory manner (Marek et al., 2018). Notably, alcohol during mid adolescence substantially disrupts mPFC dopaminergic activity, including via interactions with PV+ interneurons (Trantham-Davidson et al., 2017) and reduced D1 receptor modulation in mPFC-BLA circuits (Obray et al., 2022). Alcohol in early life also appears to reduce mPFC PV+ expression (Hamilton et al., 2017) and may restrict PV+ neuronal plasticity by increasing the density of surrounding perineuronal nets (Dannenhoffer et al., 2022).

### Concluding Remarks

Our findings support theories arguing that early-mid adolescence, which coincides with pre– and peripubertal periods (McCormick, 2022), is a particularly sensitive developmental window where alcohol exposure can pervasively impact mesolimbic and corticolimbic circuit refinement. Our results are congruent with disturbed developmental trajectories of limbic regions like BLA, mPFC, and hippocampus, and potentially including dopaminergic function as they related to behavior in ambiguous contexts. These data also extend previous work from our laboratory showing that early life adversity disturbs fear-related learning in adulthood (Bercum et al., 2021, Bercum et al., 2023).

It is difficult to extricate the degree to which the effects observed in this study were due solely to the voluntary consumption of alcohol or any stress associated with the drinking paradigm (Sicher et al., 2022). Indeed, the modified DID paradigm we used required that rats be socially isolated and denied normal drinking water during assigned drinking periods. Often after ethanol consumption, rats then satiated with water after which we also qualitatively observed high frequencies of social play bouts, especially among the Early-EtOH rats, which is a behavior frequently observed when young rodents have been exposed to stressful social isolation and denied opportunities to play with age-matched counterparts (Cooper et al., 2023). However, subsets of water-drinking rats assigned to DID conditions were, as adults, behaviorally indifferentiable from those that remained group-housed during adolescence, which suggests that any potential stress-induced effect of the DID experience likely resulted from a combination of alcohol and the nature of the modified DID approach we utilized.

It is also noteworthy that rats, especially females, have been shown to exhibit rapid activity bursts such as “darting” under fearful conditions (*e.g.* Gruene et al., 2015). While it is plausible that such rapid activity could underlie the reductions in freezing we observed during acquisition and renewal. However, automated motion detection and observations by trained experimenters confirmed that movements by in each of these conditions appeared to occur at lower velocities that do not support rapid activity bursts. In Early-EtOH rats especially, reduced freezing coincided with exploratory actions, such as high frequencies of sniffing and rearing behavior. That darting did not reliably occur for any of our subjects could be due to the small size of the conditioning chambers.

In sum, adolescence is a particularly sensitive developmental window for both sexes and perturbations during this period such as environmental toxin exposure (*e.g*. alcohol) and potentially stressful experimental paradigms (*e.g.* our modified DID paradigm) can produce lasting deficits that persist to adulthood (see Spear, 2015, Sicher et al., 2022). These findings argue for the importance of including both male and female subjects when investigating the effects of alcohol and/or stress exposure at different phases of adolescence (Sicher et al., 2022).

## Acknowledgements

This work was supported by a grant from NIDA to MPS (DA044980). Preliminary versions of this work were presented at the Stress Neurobiology Workshop and was supported by a travel award to JAG. Undergraduate research support for MV was generously provided by the Biological Science Initiative (BSI) and Undergraduate Research Opportunity Program (UROP) at CU Boulder. The authors also greatly appreciate the contributions of our many hardworking undergraduate research associates, including Richard Farrell III, Zack Marshall, Olivia Parsons, Madison Martin, Bodhi Rubenstein, Catie Bates, Nate Bosnian, G Kumar, and Makena Schutz.

